# Multiscale dynamic mean field model to relate resting-state brain dynamics with local cortical excitatory-inhibitory neurotransmitter homeostasis in health and disease

**DOI:** 10.1101/390716

**Authors:** Amit Naskar, Anirudh Vattikonda, Gustavo Deco, Dipanjan Roy, Arpan Banerjee

## Abstract

Previous neuro-computational studies have established the connection of spontaneous resting-state brain activity with “large-scale” neuronal ensembles using dynamic mean field approach and showed the impact of local excitatory−inhibitory (E−I) balance in sculpting dynamical patterns. Here, we argue that whole brain models that link multiple scales of physiological organization namely brain metabolism that governs synaptic concentrations of gamma-aminobutyric acid (GABA) and glutamate on one hand and neural field dynamics that operate on the macroscopic scale. The multiscale dynamic mean field (MDMF) model captures the synaptic gating dynamics over a cortical macrocolumn as a function of neurotransmitter kinetics. Multiple MDMF units were placed in brain locations guided by an anatomical parcellation and connected by tractography data from diffusion tensor imaging. The resulting whole-brain model generates the resting-state functional connectivity and also reveal that optimal configurations of glutamate and GABA captures the dynamic working point of the brain, that is the state of maximum metsatability as observed in BOLD signals. To demonstrate test-retest reliability we validate the observation that healthy resting brain dynamics is governed by optimal glutamate-GABA configurations using two different brain parcellations for model set-up. Furthermore, graph theoretical measures of segregation (modularity and clustering coefficient) and integration (global efficiency and characteristic path length) on the functional connectivity generated from healthy and pathological brain network studies could be explained by the MDMF model. In conclusion, the MDMF model could relate the various scales of observations from neurotransmitter concentrations to dynamics of synaptic gating to whole-brain resting-state network topology in health and disease.

## Introduction

Exploration of human brain function is undertaken at different scales, from neurotransmitter modulations that controls the gating of ion channels and gives rise to membrane potential dynamics leading to neuronal firing, which in turn get integrated across several thousands of neurons to generate characteristic spontaneous activity of brain areas. One of the most robust spontaneously generated spatiotemporal pattern of activity in brain is the resting-state activity (Raichle and Mintun, 2006; Rogers et al., 2007; Vincent et al., 2007; Mitra et al., 2014) which are observed when a person is not engaged in any specific task. Due to the simplistic nature of the stimulus condition, the resting-state activity paradigm is used as an important tool in studies of normal brain functions and also in identifying markers for various neurological disorders (Pearlson, 2017; Zhou et al, 2017). Some studies have also attempted to understand the underlying mechanisms at the systems level that govern the temporal dynamics of resting-state networks (Deco & Jirsa, 2012; Deco et al., 2014a, Demirtas et al 2017, see also Cabral et al., 2017 for a review).

All these studies use neuro-computational models as a mechanism to link the spontaneous resting-state activity to the dynamics of neuronal ensembles (more technically called the mean field activity) located at different regions of the cortex, taking into account the anatomical constraints obtained from tractography studies. Recent neuromodeling theories suggest that resting-state correlations emerge when (i) the dynamic state of the brain is close to criticality, i.e., at the edge of a bifurcation, and (ii) the excitatory inputs are balanced by local inhibitory processes maintaining the homeostasis (Deco & Jirsa, 2012; Deco et al., 2014a; Vattikonda et al., 2016). One important challenge is to understand how do the interaction among physiological parameter(s) like excitatory and inhibitory neurotransmitter concentrations at the micro-scale level shapes up the spontaneous and structured resting-state dynamics of brain. The problem is further compounded by the fact that brain parameters, such as the proportion of different neurons and neurotransmitter types (glutamate, gamma-aminobutyric acid (GABA), etc.) with their associated synaptic properties cannot be manipulated independently in living system to delineate their role in brain function (Prinz, 2008). Due to these *in vivo* practical constraints, physiologically realistic whole brain network models are favourable candidates to overcome and manipulate experimentally inaccessible parameter(s) of neuronal networks in brain. Following this line of reasoning, mean field approach, consisting of coupled stochastic differential equations could provide appropriate neuro-computational framework to study missing link between micro-scale state variables with whole-brain network dynamics (Breakspear et al., 2017). Moreover, the computational brain network modeling could be used to explore the underlying mechanisms of human brain functions, such as at the resting-state human brain operates at maximum metastability (Deco et al., 2017).

Although molecular, cellular, and electrophysiological studies of neurons could show how post-synaptic neuronal response emerges depending on neurotransmitter types, yet its relation with global scale brain activity is not well addressed (Duncan et al., 2014). GABA and glutamate are the two key inhibitory and excitatory neurotransmitters, respectively (Novotny et al., 2003), that are present all over the cortex (Duncan et al., 2014) and govern the excitatory−inhibitory (E−I) balance (Lauritzen et al., 2012). Applying statistical measures, several studies have demonstrated inhibitory neurotransmitter has significant association with blood-oxygen-level dependent (BOLD) signal in healthy human brain (Northoff et al., 2010; Raichle et al., 2009). Moreover, interaction among inhibitory neurons and excitatory neurons is thought to have a direct influence on BOLD signal (Logothetis et al., 2001). In recent past, neuro-computational studies have shown how balance in local E−I input impacts spontaneous resting-state activity of brain (Deco et al., 2014a; 2014b). Our group had already shown the importance of E−I balance in view of functional recovery, e.g., establishment of homeostasis over a re-organized brain network following a lesion in specific brain site (Vattikonda et al., 2016). As imbalance of inhibition and excitation is the key neuronal mechanism involved in neurological disorders (Chiapponi et al., 2016), the study on glutamate−GABA balance would play a key role in exploring mechanistic insights of neurological diseases and its deviation from normal brain function.

In principle, a combination of fMRI-derived BOLD signal and magnetic resonance spectroscopy (MRS) techniques can be used to find out association of neurotransmitter concentration(s) and BOLD signal vis-a-vis resting-state. However, glutamate−GABA level estimated via MRS and its association with BOLD signal remain a matter of contentious debate (Kurcyus et al., 2018). Hence, neurocomputational models such as the one presented here provides a plausible link between rich resting-state spatiotemporal dynamics of whole-brain networks with neurotransmitter concentrations. Here, we have proposed a multi-scale model that connects the scale of synaptic gating dynamics represented by kinetic equations (Destexhe et al., 1994a) of neurotransmitter concentration gradients with the scale of mean field activity of excitatory/inhibitory neural populations (Deco et al., 2014b). The neurotransmitter concentrations were set in an optimized regime where biologically relevant metric, metastability were comparable between simulated and empirical resting-state brain dynamics (Deco et al., 2017). Subsequently, our model predicts that (i) homeostasis via E−I balance could be achieved only at a specific range of neurotransmitter concentrations, and (ii) any deviations from these optimal regimes can lead to a variety of pathological conditions such as epilepsy and schizophrenia which we further verify based on multiple network measures of integration and segregation.

## Material and Methods

We propose the multiscale dynamic mean field (MDMF) model for connecting multiple scales of neuronal organization, and compare resting-state functional connectivity (rs-FC) predicted by our model guided by structural connectivity (SC) with the empirical rs-FC. We then predict global and local network measures from our model and compared them with empirical observations previously reported in literature.

### Validation on empirical data

The dataset used in this study were collected from the Cambridge Centre for Ageing, and Neuroscience (CamCAN), University of Cambridge, UK that uses 3T Siemens TIM Trio scanner with a 32-channel head coil (voxel size 3 × 3 × 4.4 mm). The SC matrix was used in the computational model to predict the rs-FC matrix. Quantitative comparison between model-generated rs-FC and empirically observed rs-FC was performed to validate our model.

### Generation of SC matrix

Empirical diffusion weighted imaging (DWI) dataset was collected from a randomized subsample of 10 healthy participants (5 males, age range 24−32; 5 females, age range 22−29) from CamCAN data set. T1 anatomical images, diffusion-weighted images, gradient vectors, and gradient values were also obtained. In this study, the empirical SC matrix was generated following seed region of interest (ROI) selection, tracking and aggregation of generated tracks by using an automated pipeline as proposed by Schirner et al., (2015). In this pipeline, high-resolution T1 anatomical images were used to create segmentation and parcellation of cortical grey matter using FreeSurfer’s recon-all function. For each subject, binary white matter (WM) masks were used to restrict tracking to WM voxels. DWI data were pre-processed using FreeSurfer following extracting gradient vectors and gradient values (known as b−table) using MRTrix. We transformed the high-resolution mask volumes from the anatomical space to the subjects diffusion space, which was used for fiber tracking. Cortical grey matter parcellation of 34 and 75 ROIs in each hemisphere was undertaken following Desikan-Killiany parcellation (Desikan et al., 2006) and Destrieux brain atlas (Destrieux et al., 2010), respectively. This was specifically performed to cross-validate the results obtained from one parcellation on the other, thus evaluating the robustness of our modelling paradigm. To estimate connection strength (value ranging from 0 to 1) between each pair of ROIs, probabilistic tractography algorithm was used to estimate how one ROI can influence the other in cortical grey matter parcellation. The pipeline aggregates generated tracks to estimate structural connectome for each individual subject. The normalized weighted distinct connection counts used here contain only distinct connections between each pair of regions yielding a symmetric matrix.

### Generation of rs-FC matrix

From the CamCAN dataset, resting-state fMRI data were used to generate rs-FC matrices employing two different parcellation schemes: 150 cortical areas (Destrieux et al., 2010) and 68 cortical areas (Desikan et al., 2006). fMRI data were collected from all participants with their eyes closed while remaining awake for 9 min 20 s. For each participant, resting-state fMRI scans were acquired at 261 time points with TR=1.97 s.

The major steps involved in generating rs-FC using CONN toolbox were as follows: motion correction, removal of non-brain tissue, high-pass temporal filtering, six-degrees of freedom linear registration to the MNI space, and extraction of BOLD signal. Each participant’s functional images were registered to pre-processed T1−anatomical images and parcellated into 68 ROIs using Desikan-Killiany brain atlas or 150 ROIs using Destrieux brain atlas. In each of the 68 or 150 ROIs, representative BOLD signal was computed by calculating mean of BOLD signals from all the voxels in the corresponding ROI. Pairwise Pearson correlation coefficients were computed among ROIs from the z-transformed BOLD time series to generate the rs-FC matrix for each subject. Thus, the generated rs-FC matrix represents correlation of the BOLD activity between brain regions.

#### Computational model

Previous studies have shown that the dynamic mean field (DMF) approach (Deco et al., 2014a; 2014b) is able to capture the rs-FC using feedback inhibition control to maintain E−I balance and constraining network dynamics with empirical SC extracted from the density of white matter fiber tracts connecting brain areas. One can go further to identify the role of local E−I homeostasis in shaping up rs-FC when structural connections are perturbed (Vattikonda et al., 2016). At the neurotransmitter level, the kinetic model of receptor binding (Destexhe et al., 1994a; 1994b) relates the neurotransmitter concentration changes to the synaptic gating variable. We propose that a multiscale model should be able to relate the neurotransmitter modulations to the rs-FC. The key link is the synaptic gating dynamics which is linked to neurotransmitter concentration kinetics and further plays a key role in generation of excitatory-inhibitory currents in a local population of neurons (mean field). We relate the mean field activity in an area to the BOLD activity by the haemodynamic model (Friston et al., 2000; 2003). Each cortical region is modeled as a pool of excitatory and inhibitory neurons with recurrent excitatory-excitatory, inhibitory-inhibitory, excitatory-inhibitory and inhibitory-excitatory connections, and coupled with neurotransmitters GABA and glutamate via NMDA (N-methyl-D-aspartate) and GABA synapses, respectively (Fig 1). Long-range connections are modeled as excitatory connections between excitatory pools of each region. Long-range inputs are also scaled according to connection strength between regions derived from diffusion tensor imaging. Here, excitatory population in an area receives following input currents: recurrent inhibitory currents, recurrent excitatory currents, long-range excitatory currents from excitatory populations in all other areas, as well as external currents.

**Fig. 1.**
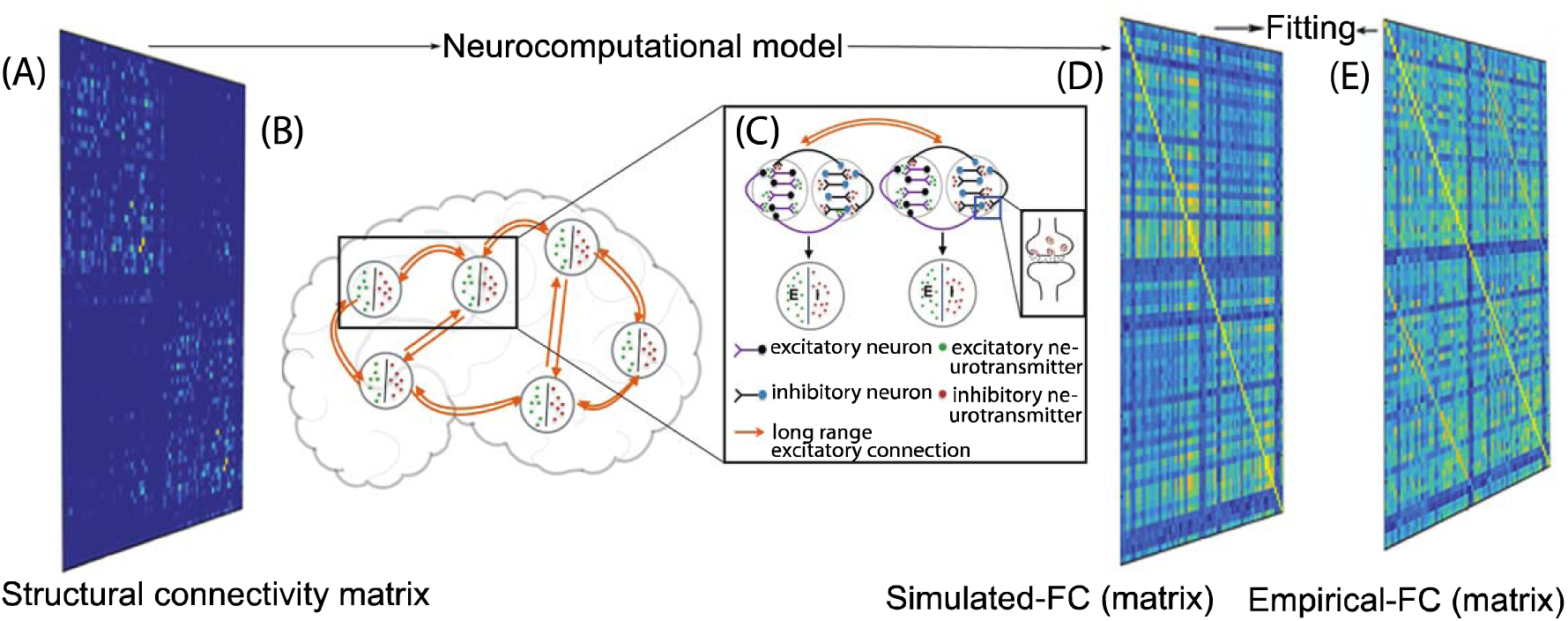
Multiscale dynamic field model (MDMF) set up (A) Averaged anatomical connectivity or structural connectivity matrix of healthy human brains. (B) Whole-brain network composed of interconnected nodes. (C) Local networks, represented by nodes or cortical areas, with each separate node consisting of excitatory (purple) and inhibitory (black) neurons with recurrent inhibitory−inhibitory, excitatory−excitatory, inhibitory−excitatory, and excitatory−inhibitory connection, and all nodes are discrete interconnected with long-range excitatory neurons (orange colour line). Each brain area is represented as a pool of excitatory (E, green) and inhibitory (I, red) neurotransmitters. In inset image, presynaptic neuron releases neurotransmitter in the synaptic cleft, and it binds with postsynaptic receptors. (D) Model generated FC matrix, and (E) its fit or similarity with empirical FC.

Destexhe et al., (1994a); assumed that occurrence of neurotransmitter release at synaptic cleft can be considered a pulse following the arrival of an action potential at the presynaptic terminal. Thus, neurotransmitter release can be captured in the rate equation that describes the synaptic gating variable dynamics. The rate of change of synaptic gating variable can be expressed as neural response function (δ-spikes) scaled by neurotransmitter concentration and the closed synaptic gating probability plus the contribution of leaky synaptic gating originating from fast processes.

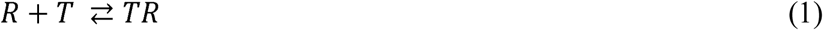

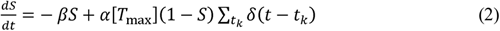

Where, *T*, *R* and *TR* represents neurotransmitter concentration, unbound and bound receptors, respectively. α and β are the forward and backward rate constants of binding of neurotransmitter on receptors. *T_max_* is maximun neurotransmitter concentration. In equation (2), *S* represents the fraction of synaptic gating variable in open state. Thus, equations (1) and (2) together describe evaluation of synaptic gating variable with time as a function of neurotransmitter concentration.

Based on the kinetic model of receptor binding (Destexhe et al., 1994a; 1994b), we have considered average release of glutamate (*T*_glu_; concentration of glutamate) or GABA (*T*_gaba_; concentration of GABA) in synaptic cleft is governed by firing rate of excitatory or inhibitory presynaptic neurons in an area *i*, and rate of synaptic gating is represented by equation (7) and equation (8), respectively.

Using DMF model (Deco et al., 2014a; 2014b) and kinetic model of receptor binding (Destexhe et al., 1994a; 1994b), we propose a MDMF model in which for each brain region the dynamics of inhibitory GABA synaptic gating and excitatory NMDA synaptic gating are governed by GABA and glutamate concentration, respectively. MDMF model is described by the following coupled nonlinear differential equations:

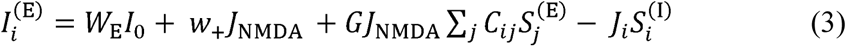

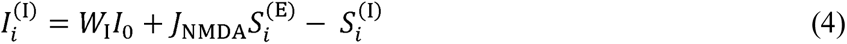

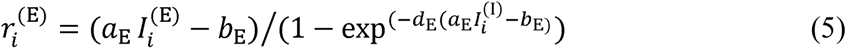

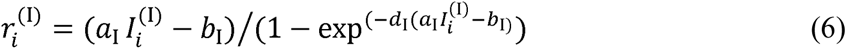

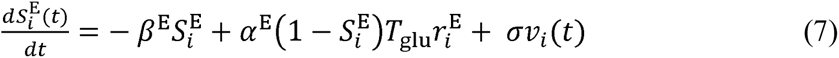

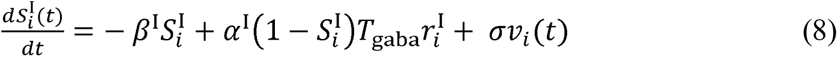

Inhibitory plasticity rule (Hellyer et al., 2016)

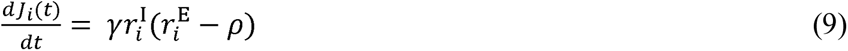

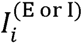, is the input current to the area *i*, where superscripts I and E represent inhibitory and excitatory populations. Firing rate of excitatory and inhibitory populations of an area are given by *r_i_*^(E)^ and *r_i_*^(I)^, respectively. *S_i_*^(E)^ and *S_i_*^(I)^ are average synaptic gating variable of area *i*. Effective external input, *IO* is scaled by *W_I_* and *W_E_* for excitatory and inhibitory populations. Excitatory synaptic coupling strength is given by *J*_NMDA_, while *J_i_* denotes synaptic coupling strength from inhibitory to excitatory subpopulation. *V_i_* in equations (7) and (8) is uncorrelated standard Gaussian noise with amplitude of *s* = 0.001 nA. Equation (9) describes the dynamics of *J_i_*, which is inhibitory plasticity rule representing changes in *J_i_* (synaptic weight) to ensure that inhibitory current clamps the excitatory current of a population and thereby maintaining a homeostasis. *G* represents global coupling strength scaling long range excitatory connections in equation (3). Homeostasis is achieved with *J_i_* dynamics such that firing rate of the excitatory population is maintained at the target-firing rate *ρ* = 3 Hz, and is the learning rate in equation (9). The chosen target-firing rate is the firing rate observed when the inhibitory and excitatory currents are matched. *C_ii_* represents anatomical connectivity strength, derived from diffusion imaging, scaling long range excitatory currents from region *j* to region *i*. Since recurrent excitatory currents are already taken into account while computing the input current to an excitatory population, all the diagonal elements *C_ii_* are set to zero in the SC matrix. Input currents to an inhibitory population in area *i* are as follows: recurrent inhibitory currents, recurrent excitatory currents, and external currents. All the parameters in model are set to same values as in Deco et al., 2014a; 2014b. Synaptic activity of each area is used as input to the hemodynamic model (Friston et al., 2000; 2003) to generate the resting-state BOLD responses of each brain area. Differential equations are solved numerically using Euler−Maruyama method with time-step of 0.1 ms and all simulations are performed in MATLAB. All the parameters used for the simulation are given in table 1. Simulations were carried out across glutamate (0.1 to 15 mmol) and GABA (0.1 to 15 mmol) concentration regimes that span the parameter space reported from healthy and diseased brains. However, to highlight an optimized range of neurotransmitter concentrations where normal resting-state brain function can be defined we undertake the following analysis.

**Table 1.** Parameters used to simulate MDMF model

|  |  |  |  |
| --- | --- | --- | --- |
| $I_0$ | 0.382 nA | $\alpha^E$ | $0.072 \text{ ms}^{-1} \text{ mM}^{-1}$ |
| $J_{\text{NMDA}}$ | 0.1 | $\beta^E$ | $0.0066 \text{ ms}^{-1}$ |
| $W_E$ | 1.0 | $\alpha^I$ | $0.53 \text{ ms}^{-1} \text{ mM}^{-1}$ |
| $W_I$ | 0.74 | $\beta^I$ | $0.18 \text{ ms}^{-1}$ |
| $w_+$ | 1.4 | $\sigma$ | 0.001 (nA) |
| $a_I$ | 615 ($\text{nC}^{-1} \text{ l}$) | $G$ | 0.69 |
| $b_I$ | 177 (Hz) | $\gamma$ | 1 |
| $d_I$ | 0.087 (s) | $\rho$ | 3 |
| $a_E$ | 310 ($\text{nC}^{-1} \text{ l}$) | $d_E$ | 0.16 (s) |
| $b_E$ | 125 (Hz) | | |

### Using the measure of metastability to define optimal parameter space

Metastability measures the tendency to deviate from stable manifolds in a dynamical systems. For functional brain network, the stable manifold may be the modes of synchronization captured by phase locking indices. Earlier research have proposed metastability is an important measure to describe information processing in a resting-state brain (Deco et al., 2017; Alderson et al., 2018). Here we define the parameter ranges of GABA/ glutamate concentrations for which metastability computed from simulated and empirical BOLD time series were maximally consonant.

In the present study, metastability is measured using standard deviation of the Kuramoto order parameter across time. Kuramoto order parameter captures the degree of synchronization among a bunch of oscillators, and is defined by the following equation:

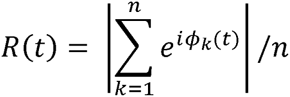

where, *Φ_k_*(*t*) denotes the phase of each narrowband BOLD signal at node *k*, and *n* is the total number of nodes. Order parameter *R(t)* measures the degree of synchronization of the *n* oscillating nodes at the global level. If *n* phases are uniformly distributed then *R* takes on value ∼ *1/n*, whereas if phases are perfectly synchronised then *R=1*. Thus, the standard deviation of *R*(*t*) estimates the tendency of the system to deviate further away from the synchronization manifold and can qualify as a measure for metastability. Standard deviation of order parameter is calculated from empirical resting-state fMRI signals and from simulated BOLD signals. First, simulated and empirical BOLD signals are processed with band-passed filter within the narrowband 0.03-0.06 Hz. Earlier findings report the narrow frequency bands have been mapped to the grey matter and it has been found to be functionally relevant compare to other frequency bands (Achard et al., 2006; Deco et al., 2017). Hilbert transform is then used to compute the instantaneous phase of each narrowband signal *k* using MATLAB function hilbert.m.

### Graph-based metrics to assess network topologies

Adjacency matrices were constructed from rs-FC for applying graph theoretical metrics in the following steps. Both negative and positive correlations are treated as connection weights by taking their absolute values. This absolute rs-FC matrix is thresholded with the correlation value of 0.25; this threshold is selected to eliminate low temporal correlation between ROIs related to signal noise (Buckner et al., 2009). Finally, thresholded rs-FC matrices were used to generate binary adjacency matrices for the various connection densities. Brain Connectivity Toolbox (BCT) is used to compute all the graph-based measures that quantifies segregation and integration in information processing among brain areas as described below.

#### Integration measures

*Global efficiency:* it quantifies efficient exchange of information across the entire network (Wang et al, 2010). Global efficiency of graph *G* is denoted by following formula: 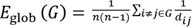

where *i* and *j* represent vertices in the graph, *n* is number of vertices, and *d_ij_* is the shortest path length between node *i* and node *j* in *G*.

*Characteristic Path Length:* average shortest path length between all pairs of nodes in the network and measures efficiency of information transfer in a network.

#### Segregation measures

*Clustering coefficient* (*C_p_*): *C_p_* of a network is defined as the average of the clustering coefficients over all nodes in the network, where the clustering coefficient *C_i_* of a node *i* is calculated using following equation.

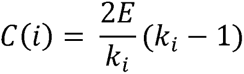

where *E* represents the number of connections among the node *i*’s neighbours and *k*_i_ is the degree of node *i*.

*Local efficiency*: it quantifies how well a network can exchange information when a node is removed.

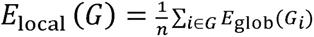

*E_glob_*(*G_i_*) is the global efficiency of sub-graph *G_i_*, which is comprised of the neighbors of the node *i*

*Modularity*: it quantifies the degree to which a network may be subdivided into clearly delineated groups. It is defined as

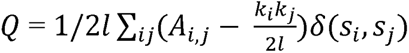

where *l* is the total number of edges, *s_i_* denotes the community to which is assigned, *δ*(*s_i_, s_j_*) is 1, if *s_i_* = *s_j_*, and 0 otherwise, *k_i_* and *k_i_* are the degree of nodes, and *A_i,j_* is the number of edges between vertices *i* and *j*.

### Data and Code Availability

All data and codes will be made available by the corresponding authors on request.

## Results

### Whole brain MDMF model predicts rs-FC

The architecture of the MDMF model consisting of cortical areas or nodes that are interconnected with structural connections are represented in Fig.1A. The whole-brain network (Fig.1B) with long-range excitatory projections among distributed brain areas contribute to resting-state brain activity. By construction the arrival of action potential in presynaptic terminal releases neurotransmitter in the synaptic cleft and it binds to the other side of cleft on receptors of post-synaptic membrane (inset image of Fig.1C). We make the assumption that released neurotransmitter at synaptic cleft appears as a pulse following arrival of an action potential at the presynaptic terminal. The total neurotransmitter release in synaptic cleft is modelled as the summation of δ-like spikes generated following arrival of action−potentials at the presynaptic terminal.

Each cortical area is described as a pool of excitatory and inhibitory neurons with recurrent excitatory−excitatory, inhibitory−inhibitory, excitatory−inhibitory, and inhibitory−excitatory connection, whereas long-range connections are modelled as excitatory connections between excitatory pools of each region (Fig.1C). Synaptic gating depends on neurotransmitter released in the synaptic cleft, hence average synaptic gating is governed with mean release of neurotransmitter in a node. So, the ensemble activity of each cortical area is the outcome of E−I neurotransmitter homeostasis (Fig.1C). MDMF uses anatomical structural connectivity matrix to connect the distributed brain regions, following which dynamic interactions equations (3)−(9) generate BOLD signals in each parcellated brain region. Finally, functional connectivity matrix computed from simulated BOLD signal is matched with empirical rs-FC (Fig.1D and E).

### Steady-state solutions of the MDMF model

In Fig.2A and B, GABA- and glutamate- mediated synaptic gating across population mean firing rate are generated by taking the steady-state solutions for equations (7) and (8) of the MDMF model, along with GABA- or glutamate-mediated synaptic gating from the model proposed by Destexhe (1994b). We use fixed values of glutamate (7.46 mmol) and GABA (1.82 mmol) concentrations observed in Precuneus of normal healthy individual’s brain during resting-state and reported in magnetic resonance spectroscopy (MRS) study by Hu et al. (2013). Fig.2 shows that in terms of average gating kinetics, the MDMF model results are equivalent to the model proposed by Destexhe (1994b), Then, we have varied *G* (global coupling strength) with fixed concentration of glutamate (7.46 mmol) and GABA (1.82 mmol) and solved the system of equations (3)−(9) numerically. Upon examining FC distance (Fig.2C), average firing rate of excitatory populations (Fig.2D) and average firing rate of inhibitory populations (Fig.2E) as a function of G, maximum similarity between simulated rs-FC and empirical rs-FC was observed at 0 = 0.69 (Fig.2C). At this 0 value, average firing rate of cortical excitatory populations is ∼4 Hz, which is within the observed range of mean firing rate of excitatory population in cortex (∼3-6 Hz; Bittner et al., 2017), and average firing rate of cortical inhibitory populations is ∼9 Hz (Fig.2E). In Fig.2F and Fig.2G, firing rate of excitatory population and inhibitory population, respectively, of each cortical area are shown in all 68 brain areas (full name of ROIs in each hemisphere is provided in table 2).

**Fig. 2.**
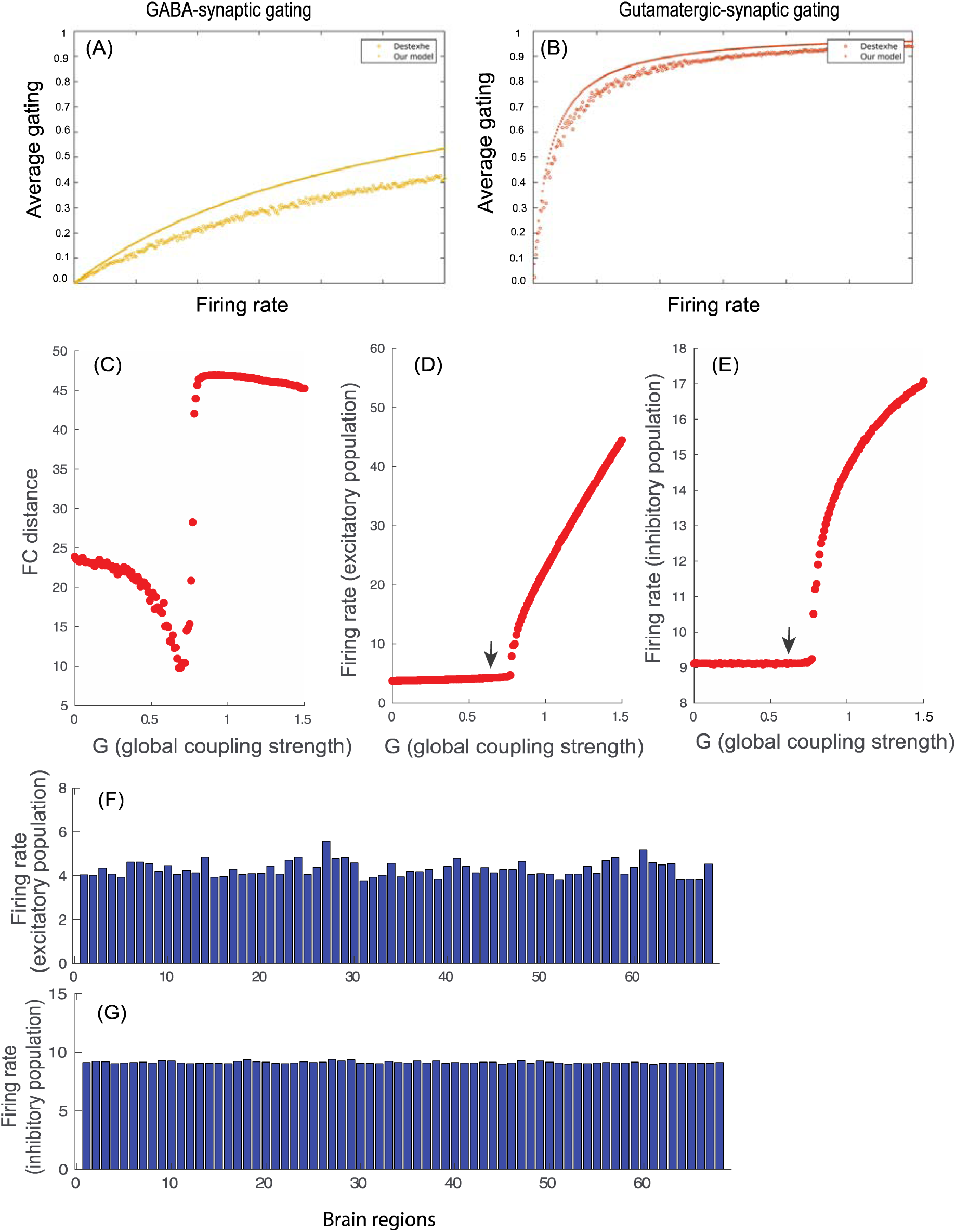
Comparison of MDMF model prediction of synaptic gating variables at steady state and the gating kinetics model proposed by Destexhe (1994b). Model-generated average (A) GABA- or (B) NMDA-synaptic gating as a function of population mean firing rate (dotted lines) closely approximate the model proposed by Destexhe (1994b) (lines with empty circles). (C) FC distance are examined as a function of G, each point represents FC distance for each G value. For G= 0.69, FC distance is found to be minimum. Average firing rate of excitatory population as a function of G, each point represents average firing rate of excitatory population for a specific G value. (D) For G=0.69, average firing rate of excitatory population is ∼4 Hz (represented by black arrow). (E) Mean firing rate of inhibitory population as a function of G, each point represents mean firing rate of inhibitory population for a specific G value. For G=0.69, average firing rate of inhibitory population is ∼9 Hz (denoted by black arrow). (F) Firing rate of excitatory populations of all 68 brain areas (Desikan-Killiany atlas) is shown for G=0.69. (G) Firing rate of inhibitory populations of all 68 cortical areas is shown for G=0.69.

**Table 2.** All 34 ROIs in each hemisphere. Area ID denotes the order of ROIs in the functional and structural connectivity matrices for each hemisphere.

| Area ID | Abbreviation | Full name |
| --- | --- | --- |
| 1. | BSTS | Banks of Superior Temporal Sulcus |
| 2. | CAC | caudalanteriorcingulate |
| 3. | CMF | Caudalmiddlefrontal |
| 4. | CUN | Cuneus |
| 5. | ENT | Entorhinal |
| 6. | FUS | Fusiform |
| 7. | IP | Inferiorparietal |
| 8. | IF | Inferiortemporal |
| 9. | IC | Isthmuscingulate |
| 10. | LO | Lateraloccipital |
| 11. | LOF | Lateralorbitofrontal |
| 12. | LIN | Lingual |
| 13. | MOF | Medialorbitofrontal |
| 14. | MT | Middletemporal |
| 15. | PH | Parahippocampal |
| 16. | PC | Paracentral |
| 17. | PAOP | Parsopercularis |
| 18. | PAOR | Parsorbitalis |
| 19. | PT | Parstriangularis |
| 20. | PEC | Pericalcarine |
| 21. | POCE | Postcentral |
| 22. | POCI | Posteriorcingulate |
| 23. | PRCE | Precentral |
| 24. | PRCU | Precuneus |
| 25. | RAC | Rostralanteriorcingulate |
| 26. | RMF | Rostralmiddlefrontal |
| 27. | SF | Superiorfrontal |
| 28. | SP | Superiorparietal |
| 29. | ST | Superiortemporal |
| 30. | SM | Supramarginal |
| 31. | FP | Frontalpole |
| 32. | TP | Temporalpole |
| 33. | TT | Transversetemporal |
| 34. | INS | Insula |

Henceforth, *G* = 0.69 is selected for all the subsequent analysis.

### Determining local homeostasis regime of GABA and glutamate concentrations relating to rs-FC

We solved the system of equations (3)−(9) numerically over a parameter space of glutamate and GABA concentrations from 0.1 to 15 mmol. Mean of firing rates of excitatory population across all parcellated cortical areas are examined at various GABA and glutamate concentrations. Two different parcellations Desikan-Killiany (Desikan et al., 2006) and Desteriux (Destrieux et al., 2010) were used to evaluate the robustness of results. Fig.3A and D, depict mean firing rate of excitatory populations across 68 and 150 parcellated cortical areas, respectively at glutamate concentration ranging from 0.1 to 15 mmol and GABA concentration ranging from 0.1 to 15 mmol. Metastability is computed using Kuramoto order parameter (see Materials and Methods section for details) from simulated BOLD signals from parcellated brain areas (Fig 3B, E) across various concentration of GABA and glutamate. Metastability computed from empirical BOLD signals obtained from subsamples of CamCAN dataset (10 subjects) is found to be ranging from ∼0.0002 to ∼0.0005. So, the region outlined by white lines in Fig.3B & E denotes parameter regimes where metastability from simulated BOLD signals matches with metastability obtained from empirical BOLD signals. Hence, using Desikan-Killiany atlas, match between metastability of simulated BOLD signal and empirical BOLD signal is found at glutamate concentration ranging from 4.4 to 9.7 mmol and GABA concentration ranging from 0.4 to 3.9 mmol (Fig.3B). Even with a finer parcellation scheme (Desteriux et al., 2010), glutamate concentration ranging from 4.2 to 9.2 mmol and GABA concentration ranging from 0.2 to 3 mmol shows good match between metastability of simulated BOLD signal and empirical BOLD signal (Fig.3E). Also the maximum similarity between the empirical rs-FC and model-predicted rs-FC, measured by the Frobenius norm is found at glutamate concentration ranging from 4.4 to 9.7 mmol and GABA concentration ranging from 0.4 to 3.9 mmol (Fig.3C). Using finer parcellation scheme (Desteriux et al., 150 parcellated brain areas) maximum similarity between the empirical rs-FC and simulated rs-FC is found at glutamate concentration ranging from 4.2 to 9.2 mmol and GABA concentration ranging from 0.2 to 3 mmol (Fig.3F). Empirical observations from MRS studies show that adult normal human brain contain glutamate from 6-12.5 mmol/kg and GABA from 1.3-1.9 mmol/kg (Govindaraju et al., 2000).

**Fig. 3.**
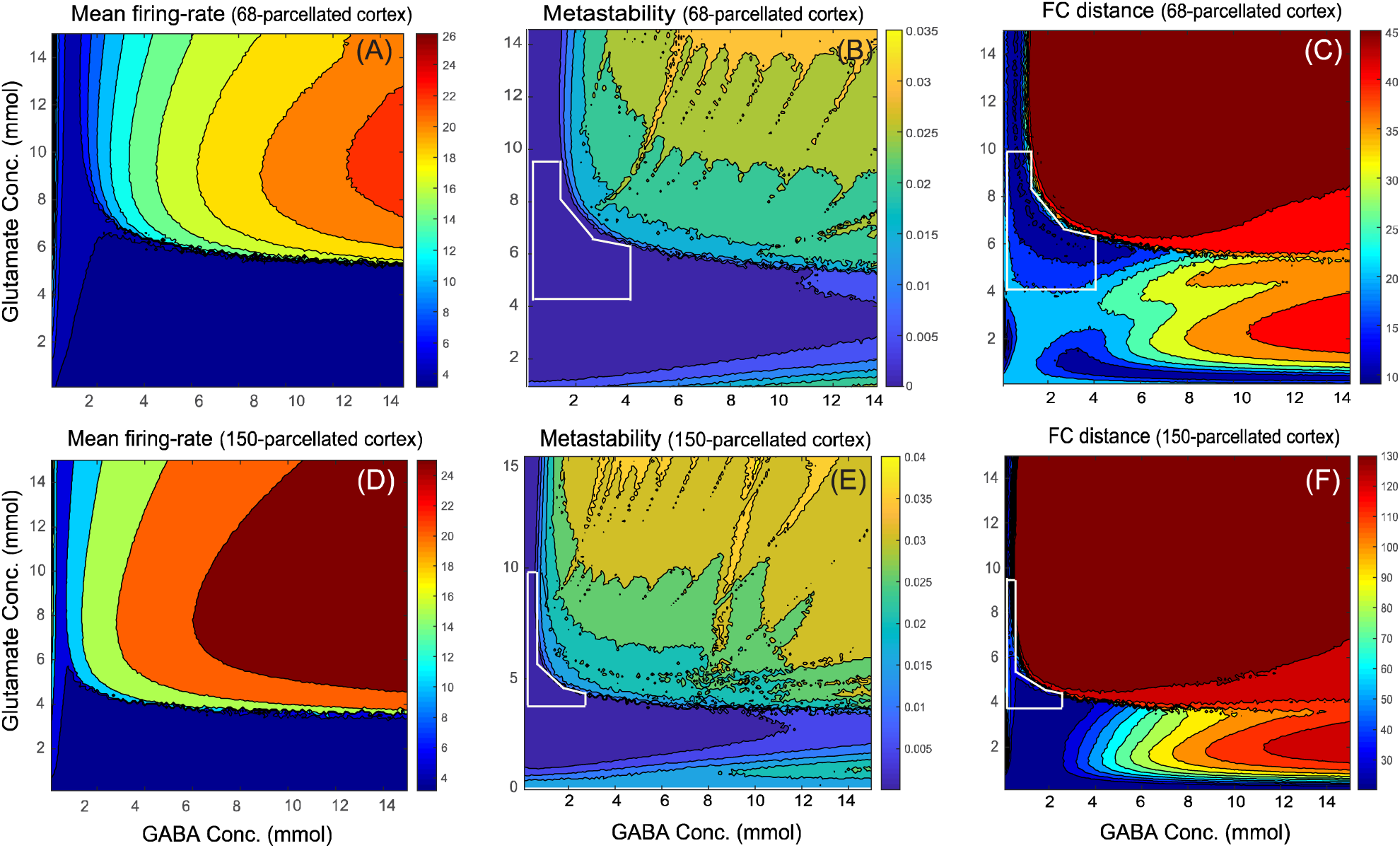
Neuronal firing, metastability, and FC distance as a function of GABA and glutamate concentrations. (A) Mean firing rate: average of firing rates of excitatory population. (B) Metastability computed from the simulated BOLD signals (C) FC Distance: similarity between empirical rs-FC and simulated rs-FC using Frobenius norm computed by solving MDMF model over various GABA (0.1−15 mmol) and glutamate (0.1−15 mmol) concentrations. One cortical unit of MDMF was chosen at each of 68 parcellated brain regions using Desikan-Killiany atlas (A, B and C) and at 150 parcellated regions using Deteriux atlas (D, E, F).

Finally, we have argue that the local homeostasis of E−I balance is obtained at glutamate concentrations ranging from 4.4 to 9.7 mmol and GABA concentrations ranging from 0.4 to 3.9 mmol using Desikan-Killiany atlas. This optimal regime of neurotransmitter concentrations is further used for simulations with finer parcellation that also yielded minimum metastability.

### Relationship between neurotransmitter concentrations and the degree of functional segregation and integration

Empirical evidences show dysfunction in glutamate-ergic and in GABA-ergic neurotransmission plays key role in epilepsy and in schizophrenia patients. The changes (increase or decrease) in neurotransmitter concentrations measured via MRS in epileptic and schizophrenia patients as compared to healthy subjects are tabulated in tables 3 and 4, respectively. The disrupted network topological properties such as segregation and integration measures of functional network involvement in epileptic (Wang et al., 2014; Yasuda et al., 2015) as well as in schizophrenia patients (Hadley et al., 2016; Ganella et al., 2017) are documented in table 5 and 6, respectively.

**Table 3.** Comparison of GABA levels measured with brain MRS recordings from epileptic patients and healthy subjects reported in the literature.

| *Sl. No.* | *Neurotransmitter* | *Epileptic patients as compared to control* | *Brain region* | *Reference* |
| --- | --- | --- | --- | --- |
| 1. | Glutamate | increased | Temporal lobe | Albrecht & Zielińska, 2017 |
| 2. | GABA | Increased | Dorsolateral prefrontal cortex | Chowdhury et al., 2015 |
| 3. | GABA | Decreased | Occipital lobe | Petroff et al., 1995 |
| 4. | GABA | Increased | Frontal area | Hattingen et al., 2014 |

**Table 4.**
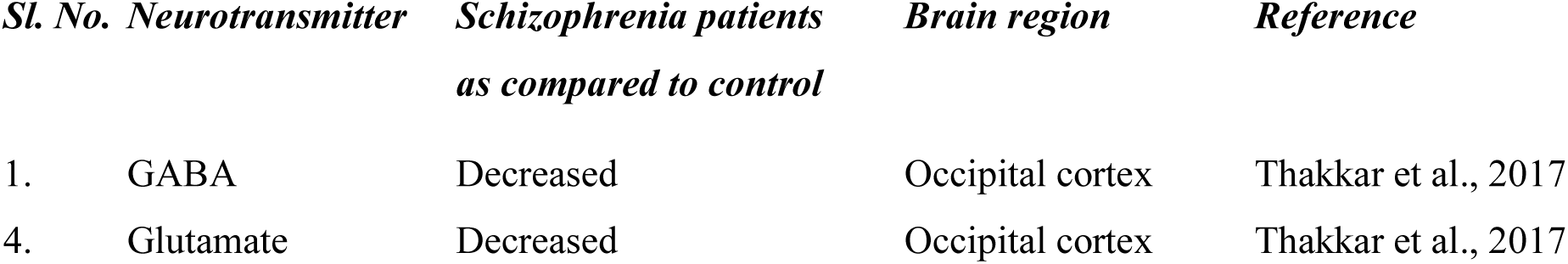
Comparison of glutamate−GABA levels measured with brain MRS recordings from schizophrenia patients and healthy subjects reported in the literature.

**Table 5.** Network segregation measures reported in literature using human brain fMRI from epileptic and schizophrenia patients relative to healthy subject discrete.

| *Sl. No.* | *Network segregation measures of brain* | *Epileptic patients as compared to control* | *Reference* | *SZ patients as compared to control* | *Reference* |
| --- | --- | --- | --- | --- | --- |
| 1. | Clustering coefficient | Increased | Yasuda et al., 2015; Jiang et al., 2017; Wang et al., 2014; Pedersen et al, 2015 | Increased | Hadley et al., 2016 |
|  | Clustering coefficient | --- | --- | Decreased | Lynall et al., 2010; Liu et al., 2008 |
| 2. | Modularity | Increased | Pedersen et al, 2015 |  |  |

**Table 6.** Network integration measures reported in literature using human brain fMRI from schizophrenia and epileptic patients relative to healthy subjects (empirical observations)

| *Sl. No.* | *Network integration measures of brain* | *Epileptic patients as compared to control* | *Reference* | *SZ patients as compared to control* | *Reference* |
| --- | --- | --- | --- | --- | --- |
| 1. | Characteristic path length | Increased | Wang et al., 2014 | Increased | Sun et al., 2016 |
| 2. | Global efficiency | Decreased | Wang et al., 2014; Yasuda et al., 2015 | Decreased | Hadley et al., 2016; Ganella et al., 2017 |

In the present study, we have computed the graph theoretical properties that quantitate functional segregation and integration from simulated functional connectivity using Brain Connectivity Toolbox (BCT). Fig.4A and B illustrate the network segregation measures, modularity and clustering coefficient, respectively, while Fig.5A and B represent network integration measures, characteristic path length and global efficiency, respectively, across various GABA and glutamate concentrations. To represent variation in graph theoretical measures at the specific neurotransmitter concentrations, Fig.4C and D represent changes in modularity and clustering coefficient, respectively, whereas Fig.5C and D illustrate variations in characteristic path length and global efficiency, respectively at the discrete concentration of glutamate (7 mmol, blue line; 9 mmol, red line) over various concentrations of GABA. Here, the homeostasis regime of GABA concentration ranging from 0.4 to 3.9 mmol are marked by shaded region. Thus, the area marked with the shaded region in the figures denotes the graph theoretic properties of healthy subjects along with the corresponding neurotransmitters (GABA and glutamate) concentration, whereas outside the marked region indicates pathological functional connectivity of human brain.

**Fig. 4.**
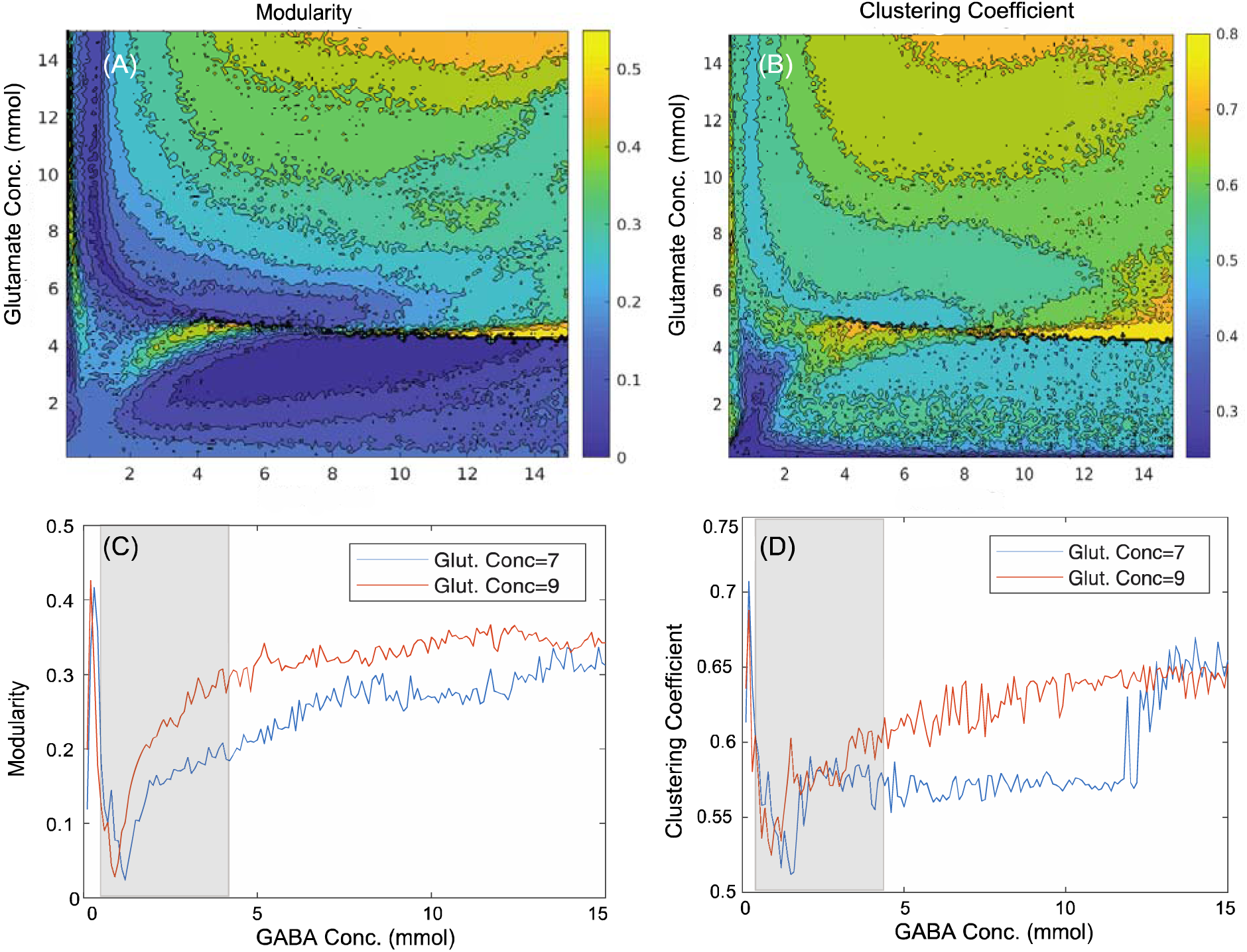
Contour plots showing the changes in network segregation measures such as, (A) modularity and (B) clustering coefficient across various concentration of GABA (0.1−15 mmol) and glutamate (0.1−15 mmol). The effect of changes in GABA concentrations ranging from 0.1−15 mmol at discrete values of glutamate concentrations (7 mmol, blue line; 9 mmol, red line) is shown on network segregation measures including (C) modularity and (D) clustering coefficient. The shaded regions denote graph theoretical measures in homeostasis regime of GABA ranging from 0.4−3.9 mmol with discrete values of glutamate (7 or 9 mmol) concentrations, whereas outside the shaded regions represent pathological scenarios.

**Fig. 5.**
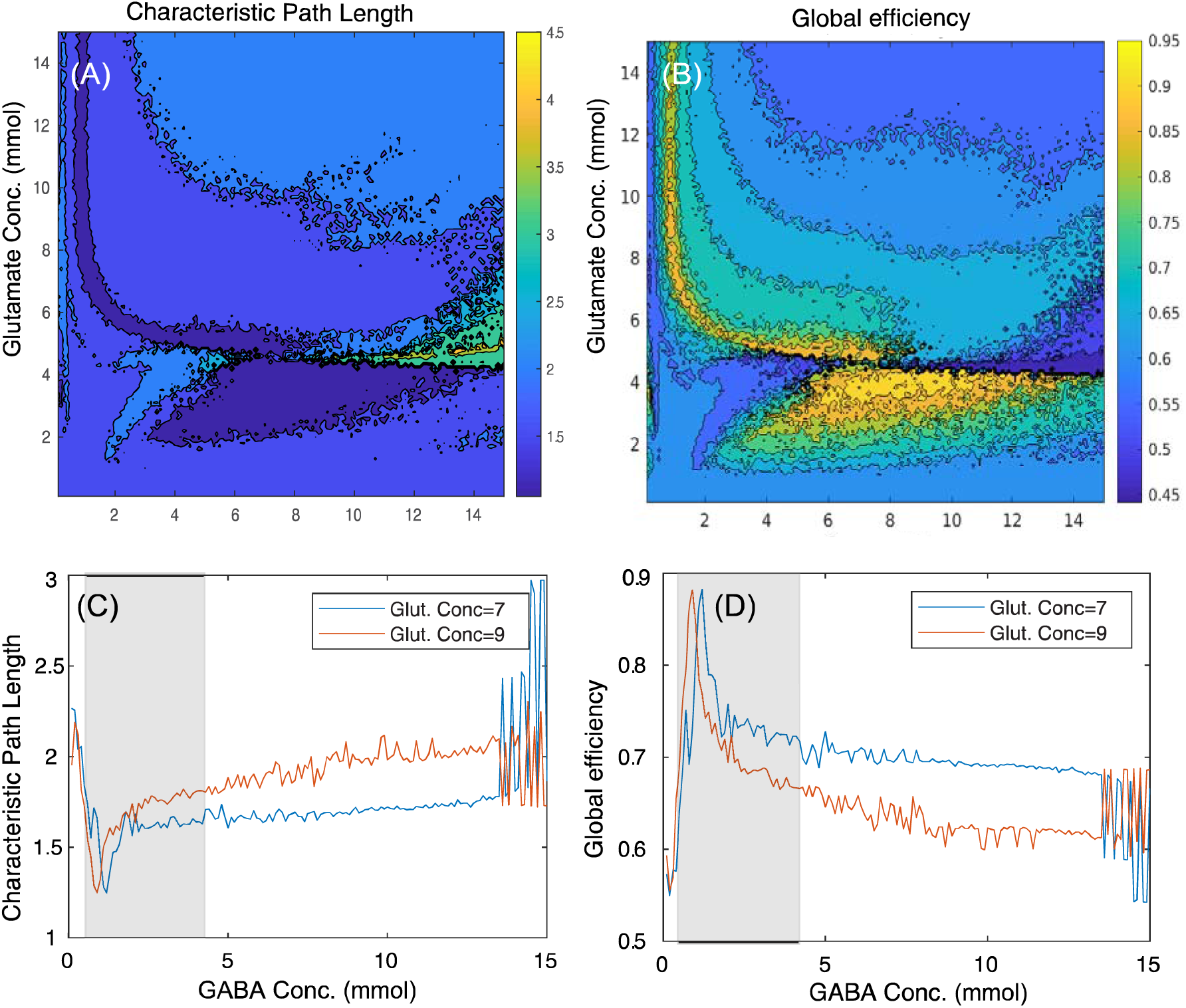
Contour plots showing the changes in network segregation measures such as, (A) modularity and (B) clustering coefficient across various concentration of GABA (0.1-15 mmol) and glutamate (0.1-15 mmol). The effect of changes in GABA concentrations ranging from 0.115 mmol at discrete values of glutamate concentrations (7 mmol, blue line; 9 mmol, red line) is shown on network segregation measures including (C) modularity and (D) clustering coefficient. The shaded regions denote graph theoretical measures in homeostasis regime of GABA ranging from 0.4−3.9 mmol with discrete values of glutamate (7 or 9 mmol) concentrations, whereas outside the shaded regions represent pathological scenarios.

The interpretations from network segregation measures (Fig.4C−D) and integration measures (Fig.5C−D) with glutamate−GABA concentrations illustrate the following scenarios of neurotransmitter level (i) low GABA concentration (0.1−0.3 mmol), (ii) homeostasis regime of GABA concentration (0.4−3.9 mmol; shaded region) and (iii) high GABA concentration (4−15 mmol) along with the discrete concentrations of glutamate (7 or 8 mmol; selected from homeostasis regime of glutamate). All the segregation measures (modularity, Fig.4C; clustering coefficient, Fig.4D) had minima for a regime of optimal GABA and glutamate values. On the other hand the characteristic path length had minima for optimal concentration of GABA−glutamate (Fig.5C), while the global efficiency peaks at the optimal concentration of GABA and glutamate (Fig.5D). An increase or decrease from the optimal concentration could be related to pathological states of the brain.

Empirical rs-FC (Fig 6A), simulated rs-FC generated at high glutamate concentration (Fig.6B), optimal glutamate concentration (Fig.6C; homeostatic regime), and low glutamate concentration (Fig.6D) with GABA concentration fixed at 1.5 mmol demonstrate that maximum similarity between empirical and simulated data (Fig.6A & C) can be achieved at optimal GABA/ Glutamate values. Euclidean distance between the simulated and empirical FC was computed using the Frobenius norm (Vattikonda et al. 2016) and was found to be minimum for optimal GABA/Glutamate (Fig 6E). In addition, FC distances computed using default mode network (DMN) nodes only, indicate high similarity between empirical and simulated data for optimal values of GABA/ Glutamate (Fig 6F). An earlier report shows that increased glutamate/GABA ratio results in enhanced correlations among DMN network nodes (Kapogiannis et al., 2013). Hence, we argue that MDMF can accurately predict the reorganization of correlations among large-scale brain networks following changes in neurochemical gradients.

**Fig. 6.**
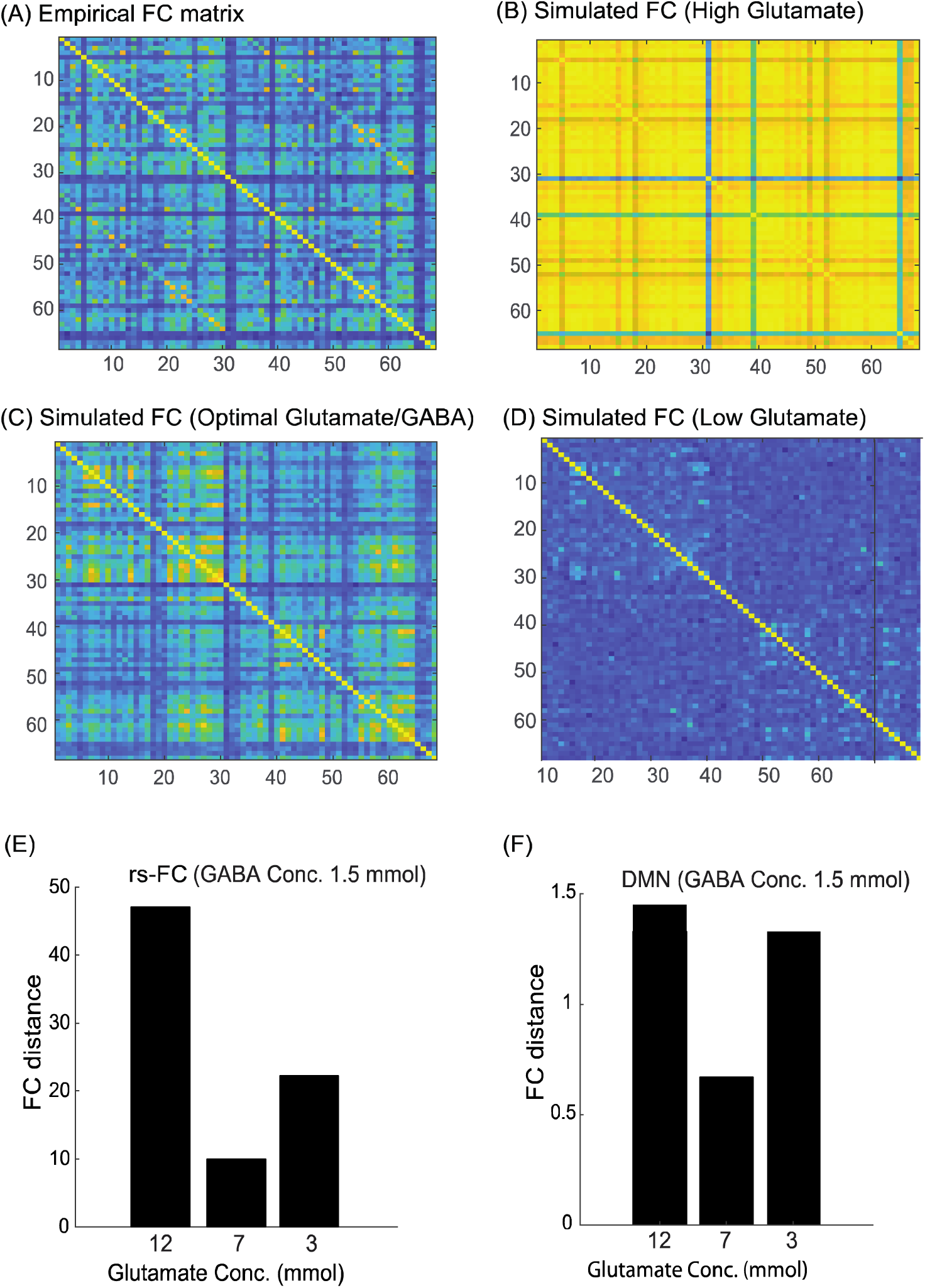
Comparison of empirical rs-FC and simulated rs-FCs. (A) Empirical rs-FC. (B) Simulated rs-FC with 12 mmol glutamate and 1.5 mmol GABA (high glutamate), (C) 7 mmol glutamate and 1.5 mmol GABA (optimal glutamate) and (D) 3 mmol glutamate and 1.5 mmol GABA (low glutamate). (E) FC-distance between empirical rs-FC and simulated rs-FC obtained at various glutamate concentrations including 12 mmol, 7 mmol or 3 mmol glutamate with GABA fixed to 1.5 mmol. (F) FC-distance between empirical DMN and simulated DMN nodes generated at different glutamate concentrations including 12 mmol, 7 mmol or 3 mmol glutamate with 1.5 mmol GABA.

## Discussion

In the present study, we have proposed a multi-scale dynamic field model (MDMF) that a provides a mechanistic explanation of whole brain resting-state network dynamics in human brain as a function of excitatory and inhibitory neurotransmitter kinetics and its interaction with the neural field. In other words, it is an effort to bridge two different scales: neurotransmitter concentration kinetics and neural field activity. MDMF brings specificity in precise quantification of the underlying neurotransmitter concentrations involved in shaping the topological properties of rs-FC, opening up possibility of tracking departures from healthy to pathological brain states in a systematic manner. The most crucial findings from the current study are (i) demonstration of a regime of optimal balance between neurotransmitter (glutamate−GABA) concentrations in the parameter space and a local homeostasis between neurotransmitter concentration and neural field (ii) graph-theoretical analysis using network segregation (modularity and clustering coefficient) and integration measures (characteristic path length and global efficiency) in the identified parameter space reconcile a wide number of contradictory empirical observations in a variety of pathological conditions.

In the MDMF model, two distinct scales of observation and measurement, the neurotransmitter kinetics and neural field dynamics contribute harmoniously to give rise to emergent functional connectivity patterns. Recent studies have shown that human brain operates at maximum metastability (Deco et al., 2017) and with the operational principles of local feedback inhibition (Deco et al., 2014a). Inhibitory plasticity rule (Hellyer et al., 2016) was employed by clamping the firing rate of cortical excitatory population at ∼3 Hz to achieve a robust parameter space for MDMF which opens up new avenues in the domain of computational neuropsychiatry. Previous studies have identified that characterization of an optimal E−I neurotransmitter homeostasis regime is critical for understanding dynamical working point shift associated with mental and neurological disorders (Cabral et al., 2012a; 2012b; Deco et al., 2014b). Thus, MDMF could be used as a computational connectomics tool by clinicians and neuroscientists apart for studying a variety of questions related to neuropsychiatric disorders. Usage of computational whole brain models in predicting seizure propagation has been recently highlighted by Proix et al, 2017. To validate the applicability of MDMF model in the diseased brain, we undertook an extensive literature research to identify the changes of glutamate−GABA concentration in neurological disorders (see tables 3 and 4).

Interestingly, deviation from optimal neurotransmitter concentration leads to prediction of pathological brain network states using MDMF. Graph properties such as modularity, clustering coefficient, global efficiency, and characteristic path length computed from simulated rs-FC, could decrease or increase depending on how far away from the optimal Gultamate-GABA concentrations are chosen. Close qualitative match between empirical reports of segregation and integration measures obtained from clinical studies and that predicted from MDMF model was achieved (see Fig 4, 5). Importantly, the MDMF model provides a global picture for the pathophysiology of neuropsychiatric and neurogenerative disorders, where the communication among brain areas are classified in terms of local and global measures, and their relationship with the underlying physiological mechanism at the molecular level. There exist couple of dissonant findings in epilepsy research that uses MRS measurements of GABA and glutamate. On one hand, some reports show decreased GABA levels (Petroff et al., 1995), whereas on the other hand some reports reveal increased GABA level (Chowdhury et al., 2015; Hattingen et al., 2014) in epilepsy patients as compared to healthy subjects. Concurrently, network segregation measures (modularity and clustering coefficient; Pedersen et al., 2015; Yasuda et al., 2015) are reported to be increased in epileptic patients. However, in case of schizophrenia, studies report increased clustering coefficient (Hadley et al., 2016) whereas reported decrease in clustering coefficient is also observed (Lynall et al., 2010), when compared with healthy subjects. Moreover the integration measure such as characteristic path length is reported to be increased whereas global efficiency is shown to be decreased in epileptic and in schizophrenia patients as compared to healthy volunteers (see tables 3 & 4). Thus, MDMF model could successfully predict the increase in network segregation and integration measure in both low or high glutamate levels, which is in accordance to empirical observations of epilepsy and schizophrenia. We argue such predictive power from a biologically realistic model has immense potential to explain inter-individual variability of metrics evaluated over large population cohorts of neurological and mental disorders as well as healthy ageing.

The present model differs substantially in many ways from previously proposed DMF model (Deco et al., 2014a) that introduced the local excitatory−inhibitory balance. In addition to being a truly multi-scale model where the synaptic gating are allowed to have dynamic properties, in MDMF model, inhibitory plasticity rule replaces feedback inhibition control (FIC) compared to the earlier models (Deco et al., 2014a; Vattikonda et al., 2016) regulating homeostatic E−I balance mechanisms, while operating in a realistic neurotransmitters concentration regimes. Our results demonstrate that even if inhibitory plasticity rule is applied locally, it can affect globally, the large-scale brain dynamics. The local inhibitory plasticity rule in the whole-brain network is biologically relevant for providing stabilization in a plastic network and regulating optimal information flow and noise correlation (Sprekeler et al, 2007; Vogels et al., 2011).

The important limitations in MDMF at the current stage is the non-incorporation of some finer details that are relevant for functional brain network dynamics. First, we have taken a fixed value of glutamate−GABA concentration over the entire brain commensurate with measurements from MRS using single-voxel spectroscopy. However, the concentrations of these neurotransmitters are variable across different areas which can be measured from positron emission tomography (PET) recordings (D’Hulst et al., 2015). Nonetheless, a detailed analysis with glutamate−GABA values in individual brain areas contributing to resting-state dynamics remains out of the scope at this point due to lack of availability of detailed data but will be an interesting issue to resolve in future as wand when such data is available. Second, communication and synchronization between brain areas are typically modulated by distant dependent conduction delays. To keep our findings tractable and to avoid complexity, delays are neglected in this model, which may serve as a critical ingredient for shaping up global brain dynamics, giving rise to phenomenon such as oscillations and other complex spatiotemporal patterns such as chaos and multi-stability (Ghosh et al., 2008; Deco et al., 2009). Third, we have avoided the incorporation of neuromodulatory effects in the MDMF model because the discussion is purely limited to relatively small time window of resting-state dynamics. However, incorporation of all these features are possible in the MDMF framework in future. In fact, the DMF component can be replaced with thalamocortical models (e.g. van Alabada et al., 2009; Freyer et al., 2011) to address homeostasis in EEG/ MEG data in future computational studies.

In summary, we have characterized the homeostasis of glutamate−GABA concentration via a realistic large-scale neural model and explained the shift of the balance reported by empirical recordings of healthy and diseased brain. The MDMF model could link multiple scales of observations from neurotransmitter concentration kinetics to neural field dynamics of resting-state brain in health and pathological observations in disease. Interestingly, the identification of the pathological neurotransmitter concentration space also opens up novel future direction in the quest of identifying specific sets of biomarkers for characterizing progression from health to disease. Another potentially interesting direction for this approach could be fMRI data during specific sensorimotor and cognitive tasks that is far less traversed at this point. Another future direction of MDMF could be geared more towards generating specific predictions during task conditions or perturbation with brain stimulations, e.g., transcranial direct stimulation/ transcranial magnetic stimulation (tDCS/ TMS) with a high degree of patient specificity. Virtual lesions can be introduced as outlined in Vattikonda et al, (2016) for identification of reorganization in the whole brain connectome as a function of neurotransmitter homeostasis. This remains a target for our future research.

## Acknowledgements

This study was supported by National Post-Doctoral Fellowship (PDF/2016/000378) from Science and Engineering Research Board (Department of Science and Technology, Government of India) to AN, NBRC Core funds, Ramalingaswami Fellowships (Department of Biotechnogy, Government of India) to DR (BT/RLF/Re-entry/07/2014) and AB (BT/RLF/Re-entry/31/2011) and Innovative Young Biotechnologist Award (IYBA) to AB (BT/07/IYBA/2013). AB also acknowledges the support of grant number F.NO.K-15015/42/2018/SP-V from Ministry of Youth Affairs and Sports, Government of India. DR was also supported by SR/CSRI/21/2016 extramural grant from the Department of Science and Technology (DST) Ministry of Science and Technology, Government of India.

